# A Novel congenic Mouse Model to study Obesity, Diabetes and NAFLD

**DOI:** 10.1101/2022.02.08.479645

**Authors:** MJ Mahesh Kumar, Shailendra Arindkar, Perumal Nagarajan

## Abstract

The genetic background of the mutant mouse strain plays an important role in disease development. We investigated whether *Lepr* mutation on different genetic background (WSB/EiJ) influences the development of obesity, diabetes and non-alcoholic fatty liver diseases (NAFLD). For this study heterozygote (*db*/+) mice were backcrossed with WSB/EiJ strain (that is resistant to obesity and NAFLD) for more than 10 generations and intercrossed for five generations having *Lepr* mutated gene incorporated in WSB/EiJ background. The newly derived congenic strain was assessed and related with the B6.db strain and WSB/EiJ strain for the development of obesity, NAFLD. The *Lepr* mutated congenic strain on the WSB/EiJ background gained significant body weight and showed other characteristic features of obesity. However, the incidence of diabetes, NAFLD, and other kidney complications were not observed in this strain. The novel congenic mouse strain having *Lepr* gene on WSB/EiJ background would be a promising model for the study of obesity without NAFLD and will also help to understand the various mechanisms involved in the development of NAFLD.

## Introduction

Obesity is a risk factor that acts as a percussor of many diseases including type-2 diabetes and non-alcoholic fatty liver disease (NAFLD). Harmful lipids accumulate in ectopic tissues and cause local inflammation that contributes to the development of endothelial dysfunction, non-alcoholic fatty liver disease, and insulin resistance ^1–3^. Besides the role of fatty acids and cytokines, and altered secretion of adiponectin also contributes to metabolic impairments in other tissues such as the liver or pancreas ^4^. There exists strong evidence for the causal link between obesity and diabetes and the effect of the influence of obesity on type 2 diabetes risk is determined not only by the degree of obesity but also by the location in the body where fat accumulates ^5^ and synergetic effect of multiple genes for disease susceptibility ^6^. The *ob/ob* and *db/db* mice are good models to understand the pathophysiology and associated metabolic complications of type-2 diabetes and obesity and is widely used. The phenotypic manifestation of these mice strains (*ob/ob* and *db/db*) is due to defect in either leptin or leptin receptor functions, respectively. Leptin, an adipokine produced by mature adipocytes plays an important role in the regulation of energy homeostasis, lipid and glucose metabolism, and immune response. Leptin deficient ob/ob mice are maintained on a C57BL/6J genetic background, and the Leptin receptor *db/db* mice are maintained on a C57BLKS/J genetic background. This difference in background imparts the phenotypic differences of severe obesity (*ob/ob)* versus severe diabetes (*db/db*) ^7^. Most of the experiments carried out on *db/db* mice are maintained on an inbred C57BL/6J genetic background. Obese B6-db exhibit mildly elevated glucose levels, hyperinsulinemia, and pancreatic islet hypertrophy, whereas obese BKS-db mice develop severe diabetes, reduced insulin levels, and islet atrophy. The genetic studies also suggested that diabetes susceptibility in these strains is under multigenic control ^7,8^. Hence background of the mice with the mutation plays an important role in obesity and diabetes in *db/db* and *ob/ob* mice.

On the other hand, the wild-derived inbred mouse strains WSB/EiJ(WSB) were inbred for over 80 generations and are genetically unique when compared with the commonly used mouse strains ^9,10^. An interesting feature of the wild-derived WSB/EiJ strain is that it is resistant to diet-induced obesity as compared to C57BL/6J ^9,10^. When this strain is fed with a high-fat diet for over 6 months, they do not gain bodyweight or any complications such as insulin resistance and NAFLD ^9,10.^ From the above studies, it was reported that this strain is lean, long-lived, and resistant to metabolic syndrome.

In the present study, we developed a congenic mouse strain using WSB mouse as a background strain and further evaluated it for the incidence of obesity, diabetes, and NAFLD. The new congenic strain could help as an additional model for obesity diabetes and NAFLD.

## Materials and Methods

### Animals

B6.BKS(D)-Lepr^db^/J (B6-db) heterozygotes and WSB /EiJ (WSB)mouse strains (Jackson Laboratory, Bar Harbor, ME) bred at the Small Animal Facility of the National Institute of Immunology were used for the study and all the procedures were approved by Institutional Animal Ethics Committee in compliance with the Committee for the Purpose of Control and Supervision of Experiments on Animals India (CPCSEA) and ARRIVE guidelines. The WSB-db congenic strain was developed by backcrossing the *Lepr db* allele from strain with B6-db with WSB strain for a minimum of 10 generations and the littermates were intercrossed for more than five generations and the male homozygous offspring were evaluated for this study. The heterozygous *db/+* animals were identified using the PCR method ^11^ and backcrossing of the heterozygotes was selected based on the genotyping results. (Supplementary Fig -1). The animals were maintained in an individually ventilated caging system on a 12: 12-h light: dark cycle and were fed with a standard pellet diet (altromin 1314 diet) with autoclaved drinking water.

### Analysis of Physiological and Biochemical parameters

Body weights measurements were taken at every 4 weeks, and the liver weights at the end of 8 weeks. The daily food intake by WSB, WSB-db, and B6-db mice strains was measured during a 7 days period. Blood was collected at the end of 8 weeks study by cardiac puncture under anesthesia (80 and 10 mg/kg body weight of ketamine hydrochloride and xylazine respectively) for various experiments mentioned below and euthanized. Serum biochemistry was performed using in-house serum auto-analyzer Screen Master 3000, Tulip, Alto Santa Cruz, India, according to the manufacturer’s instruction. For the Glucose tolerance test, the mice were fasted overnight and were injected intraperitoneally with 2 g/kg dextrose after which, the. blood glucose was measured at 0, 30, 60 and 120 min directly using a glucometer (Accu chek - Mumbai, India) using a drop of blood from the tail vein, 48 hours before euthanasia. For measuring glycated hemoglobin (HbA1c), 5 μl blood was collected from tail bleed and measured by using HbA1c Nycocard 24T kit according to manufacturer’s protocol and monitored in Nyco Card reader II (Alere Technologies AS, Kjelsasveien, Norway

### Histological analysis

The liver, kidney, pancreas, and adipose tissues were collected at the end of 8 weeks. The tissues were fixed overnight in 10% Neutral buffered formalin processed and embedded in paraffin blocks. For Hematoxylin-Eosin (H-E), masson trichrome, and picrosirus red strain tissue sections of 4-micron thickness and stained according to standard protocols. For Oil Red O staining frozen sections were used. All representative photomicrographs were captured at 200X magnifications. Immunostaining of the pancreas was performed using a primary anti-insulin antibody (Catalog No-05-1066, Millipore, USA), followed by secondary anti-mouse Alexa 594 (1:800 dilution). The confocal imaging of the sections was performed with Leica SP5 confocal microscope and the image was analyzed using LAS X confocal software (Leica, Germany).

### Quantitative PCR analysis

RNA was isolated using Trizol reagent (MRC Inc, Cincinnati, OH, USA), and cDNA was prepared according to the manufacturer’s instruction (Bio-Rad, Hercules, CA, USA). Quantitative real-time PCR was performed with SYBR green master mix (Thermo Fisher Scientific, Waltham, MA, USA) on the Master cycler RealPlex4 platform (Eppendorf, Germany). The primer pairs and PCR conditions used are listed in Table 1. 18S rRNA was used to normalize the expression of genes, and their relative expression was calculated using the 2^-ΔΔct^ method.

### Flow Cytometric analysis of immune cells in Peripheral Blood

Blood was drawn from the retro-orbital plexus of anesthetized mice in tubes containing phosphate buffer saline having EDTA. RBCs were lysed with ACK lysis buffer (Bio Legend) and cells were stained with different antibody cocktails having combinations of Anti-mouse CD3-PE, Anti-mouse B220 FITC, Anti-mouse CD49b-APC, CD4-FITC and biotinylated Anti-mouse CD8 (all from Bio legend). Streptavidin APC (BD) was employed as a secondary antibody to detect biotin-conjugated primary antibodies. 50μl of cell mixture was mixed with 50μl of antibody cocktail and incubated at room temperature for 20 min. After primary antibody incubation cells were washed and then kept in streptavidin APC secondary antibody for another 20 min at room temperature. Appropriate single-color controls were included in the analysis for compensation purposes. Data acquisition was done on FACS Canto II (BD) flow cytometer and data were analyzed with Flow Jo software (Tree Star San Carlos, CA). The proportion of T cells (CD3^+^), CD4 T cells (CD3^+^ CD4^+^), CD8 T cells (CD3^+^ CD8^+^), B220^+^ and CD49b^+^ cells were evaluated in peripheral blood.

### Statistics

The results are presented as Mean±SD. Differences between the three groups was tested using one-way and two-way ANOVA with Bonferroni post-hoc tests using GraphPad Prism (Version5.04). The value of P<0.05 was considered significant

## 3 Results

### Physiological parameters of WSB-db mice

Both B6-db and WSB-db gained significant bodyweight and liver weight respectively as compared to WSB mice However body weight gain in WSB-db mice was less as compared to B6-db mice. Few animals were monitored up to 48 weeks and B6-db and WSB-db mice had a steady increase in body weights reaching a maximum weight of 51 grams and 43 grams respectively (Fig 1 B, D). The visceral and abdominal adipose tissue in WSB-db mice was less as compared to B6.db mice (Fig1A). The average food intake of WSB, WSB-db, and B6-db mice were 3 gm/ day, 6 gm, and 8 gm respectively (Fig1 C).

**Figure 1:**
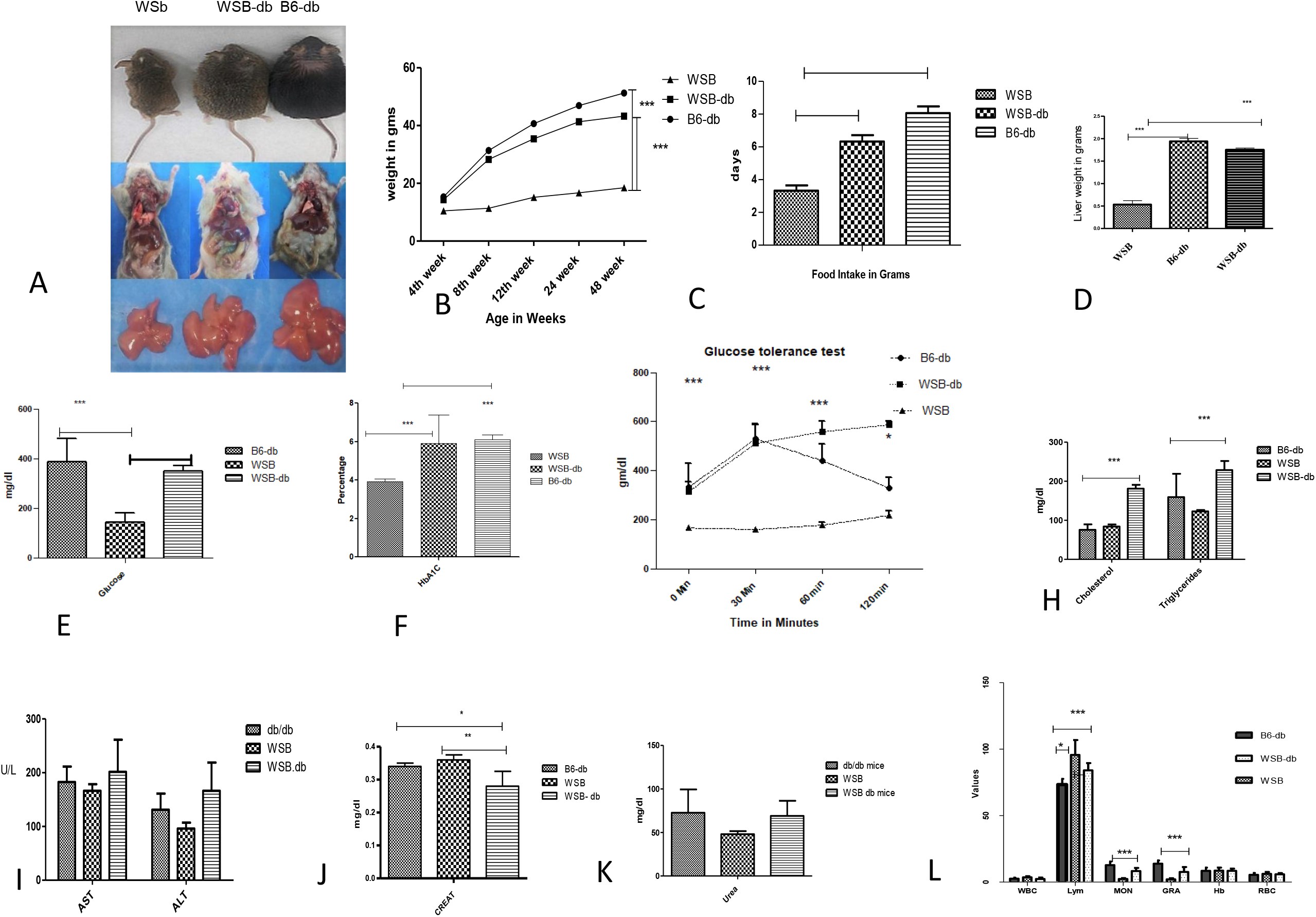
(A) Representative image showing a picture of, WSB-db, WSB and B6-db with fat deposits and liver in gross (B) represent the development of body weight gain from 4 weeks to 48 weeks in WSB-db, B6-db, and WSB mice representing a steady increase in weight gain in WSB-db mice up to 48 weeks n = 5, two-way ANOVA.(C) Mean daily food intake by WSB, WSB-db, and B6-db mice measured during a 7 days period n= 5 one-way ANOVA. (D). Mean liver weight in grams at the end of 8 weeks (n=5 one-way ANOVA). (E -F) Mean blood glucose and HbA1c level of three strains n=5 one-way ANOVA. (G) Intraperitoneal glucose tolerance test (GTT) at 8 weeks of age n = 5 2-way ANOVA), data shown as mean ± SD, *P < 0.05, **P < 0.01, ***P < 0.001, (H, I J K). Graphical representation for evaluation of serum biochemistry (CHO, TGY, ALT, AST, Creatinine, and UREA of three strains. Statistical analysis was done using 2-way ANOVA. The values are expressed in mean ± SD. *P < .05, **P < .01, ***P < .001.

### Hematology and clinical chemistry in WSB-db mice

The mean basal blood glucose, HbA1c was lesser in WSB-db as compared to B6-db mice (Fig1E, F). Both B6 Db and WSB-db strain had impaired glucose tolerance. (Fig-1G). The cholesterol and triglycerides were significantly higher in WSB-db mice as compared to WSB and B6.db mice (Fig1H). There were no significant changes in liver enzymes among these three genotypes (Fig1 I). However, the level of creatinine was significantly decreased in WSB-db as compared to WSB and B6.db mice (Fig-1 K).WSB-db mice had a significant increase in lymphocytes count with a decrease in monocytes and granulocytes count than WSB and B6.db mice (Fig-1 L)

### Histopathology and Immunofluorescence in WSB-db mice

The liver tissue of B6-db mice had ballooning of hepatocytes along with micro and macrovesicular fatty degeneration confirming steatosis only in B6.db mice (Fig 2A-C). Oil Red O-stained liver sections showed reduced lipid droplets in WSB-db mice (Fig 2D-F) and there was no marked collagen deposition in the WSB-db strain (Fig 2G-I). The kidney parenchyma of both WSB and WSB-db had normal morphology of glomerulus and tubules but there was mild pigmentation of tubular epithelial cells in B6-db mice (Fig 2J-L). The adipose tissue of WSB mice exposed normal morphology. However, the adipose tissue of B6.db and WSB-db mice had few ring-shaped adipocytes with infiltration of lymphocytes at the edges of adipocytes in B6-db mice (Fig 2P-R.). The pancreas of WSB mice had normal beta and acinar cells. WSB-db mice pancreas had moderate hypertrophy of islets with hyperplasia of beta cells. However, in B6.db mice, the pancreas exhibited moderate degenerative changes along with apoptosis of beta cells of islets (Fig 3A-C). Immunostaining of islets had more expression for antibodies for Insulin in WSB and WSB-db mice as compared to B6-db mice (Fig 3D -F).

**Figure -2.**
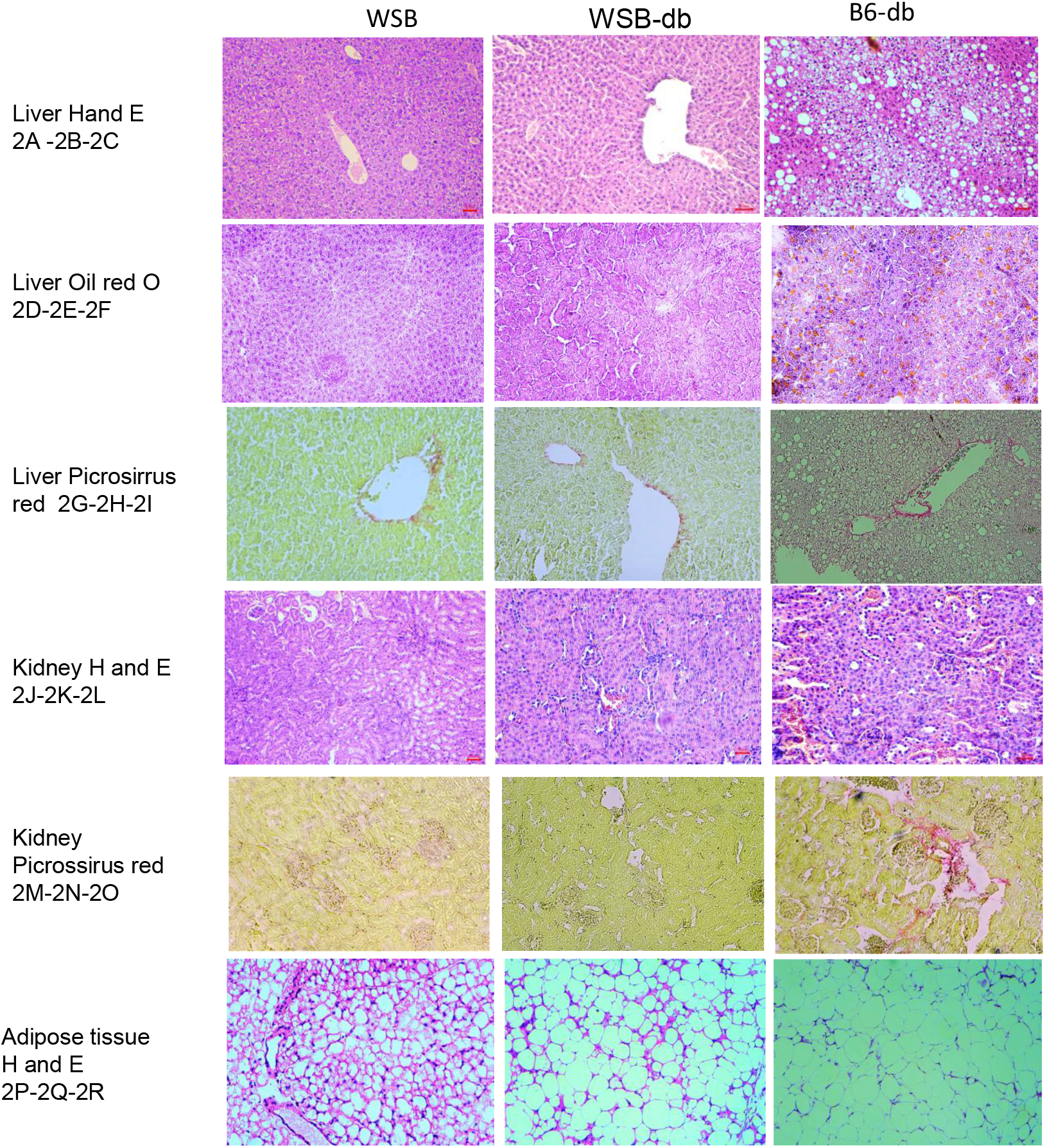
**Liver H & E: 2 A and 2 B:** hepatocytes; **2 C:** Ballooning degeneration along with macro and microvesicular fatty degeneration of hepatocytes: **Liver Oil O Red: 2 D and 2 E**: No lipid droplets observed in hepatocytes; **2 F:** Lipid droplets observed as a lobular pattern in hepatocytes : **Liver Picrosirrus red: 2 G and 2 H:** No fibrosis seen in the portal/ periportal region; **2 I**: Mild periportal fibrosis was observed : **Kidney 2H & E 2 J, K, and L**: Glomerulus and tubules appear normal: **Kidney: Picrosirrus Red: 2 M, N**: No fibrosis observed in glomerulus and tubules; **2 O:** Mild perivascular fibrosis was observed : **adipose tissue: 2 Q and R:** Few ring-shaped adipocytes with infiltration of lymphocytes at the edges.

**Figure 3.**
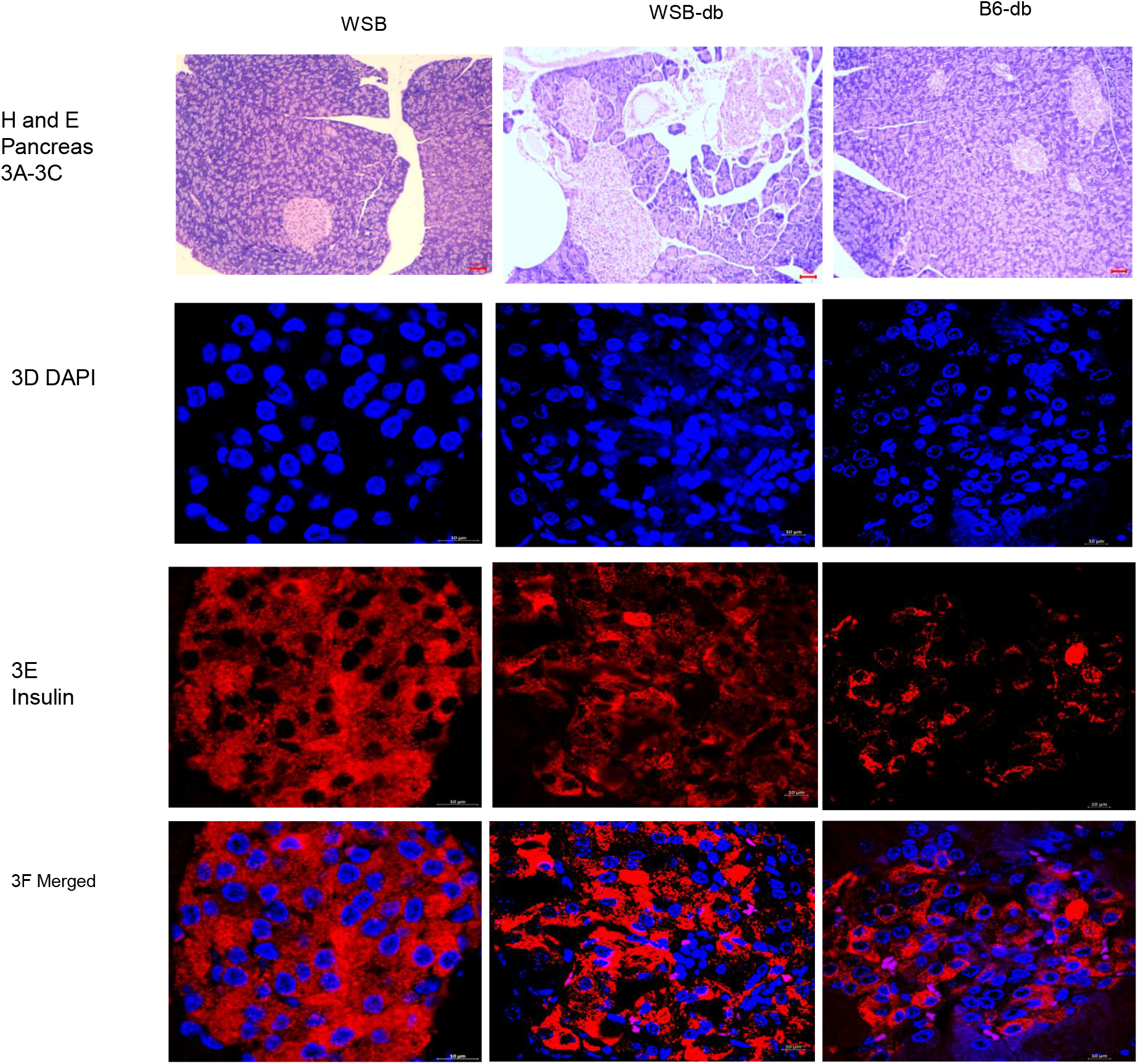
**A:** Pancreas: Normal beta cells in islets of the pancreas [Red arrow] and acinar cells in the exocrine pancreas:**3 B:** Moderate hypertrophy of islets along with hyperplasia of bet cells were observed : **3 C**: Moderate degenerative change of islets with atrophy and apoptosis of beta cells :**3D-F** Immunofluorescence; Pancreas: Merged FITC/DAPI: Overexpression of insulin protein observed in WSB-db mice compared with B6-db mice

### Gene expression analysis by reverse transcription-polymerase chain reaction

On analysis of the relative messenger RNA (mRNA) expressions of various genes involved in *denovo lipogenesis*, energy metabolism, inflammation, and insulin resistance genes in the liver of B6.db WSB-db WSB mice, the levels of denovo lipogenesis genes (ChREBP, SREBP-1c, FAS, and SCD 1) energy metabolism genes (PGC1a, PGC 1c, and PPAR alpha) genes involved in inflammation (CRP, MCP-1, CPT1a) the level of expression were higher in B6.db mice as compared to WSB-db and WSB mice. Interestingly the expression of genes related to insulin resistance (IRS-1 and IRS -2) the expression level of IRS-1 and IRS -2 in WSB mice was higher as compared to B6-db mice and WSB-db mice (Fig-4A-D)

### Flow cytometric Analysis

To investigate the immune cell changes we analyzed the level of CD3 ^+^, CD4 ^+^, CD8 ^+^, B220^+^ and CD49b ^+^ cells in peripheral blood mononuclear cells (PBMCs). We observed that there were no significant changes in the number of CD3, CD4, CD8, and CD49b cells in the peripheral blood among B6-db, WSB, and WSB-db mice. However, there was a significant increase in B220 ^+^ cells in B6-db mice as compared to WSB-db mice [Fig 5].

**Figure -4.**
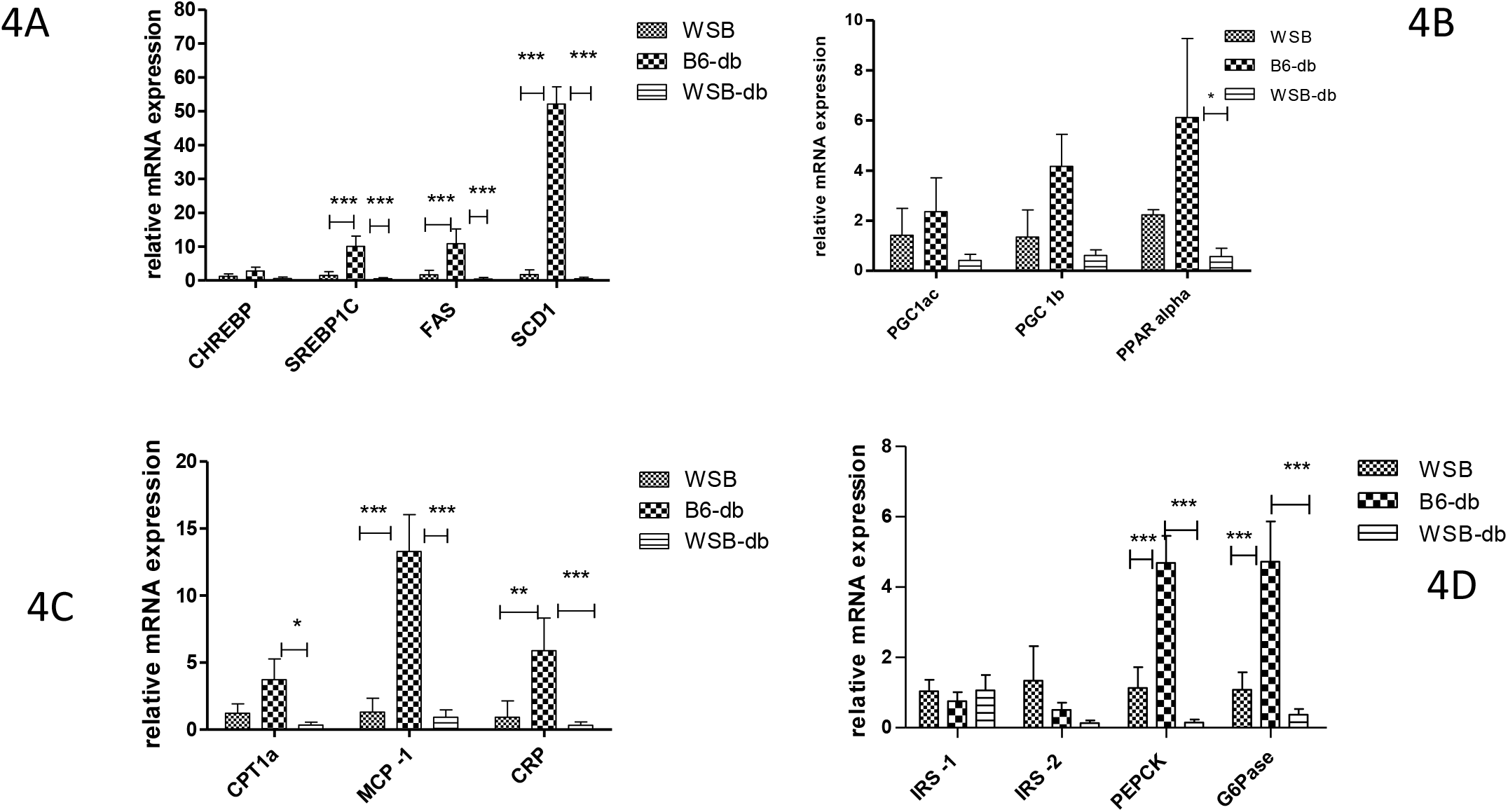
The level of expression of different genes. (A) Genes related to *denovo lipogenesis* (FAS, SCD, and SREBP1c) had significant up-regulation in B6-db mice as compared to WSB and WSB-db mice. (B) The genes involved in energy metabolism (PGC1a, PGC 1c, and PPAR alpha) exhibited up-regulation in B6.db mice as compared to WSB-db and WSB mice. (C) Represents the expression levels of genes involved in inflammation (CRP, MCP-1, and CPT1a) that were higher in B6-db mice as compared to WSB-db and WSB mice. (D) represents the expression of genes related to insulin resistance the expression level of IRS-1 and IRS -2 in WSB mice was higher as compared to B6.db mice and WSB-db mice. To compare the expression of mRNA liver samples from three animals were extracted followed by cDNA synthesis and qRT-PCR was set up in triplicate. The average Ct values of these triplicate samples were used for the calculation of fold change expression for different genes. Two-way ANOVA. test was performed and the values are expressed in mean ± SD *P < .05, **P < .01, ***P < .001.

**Figure -5.**
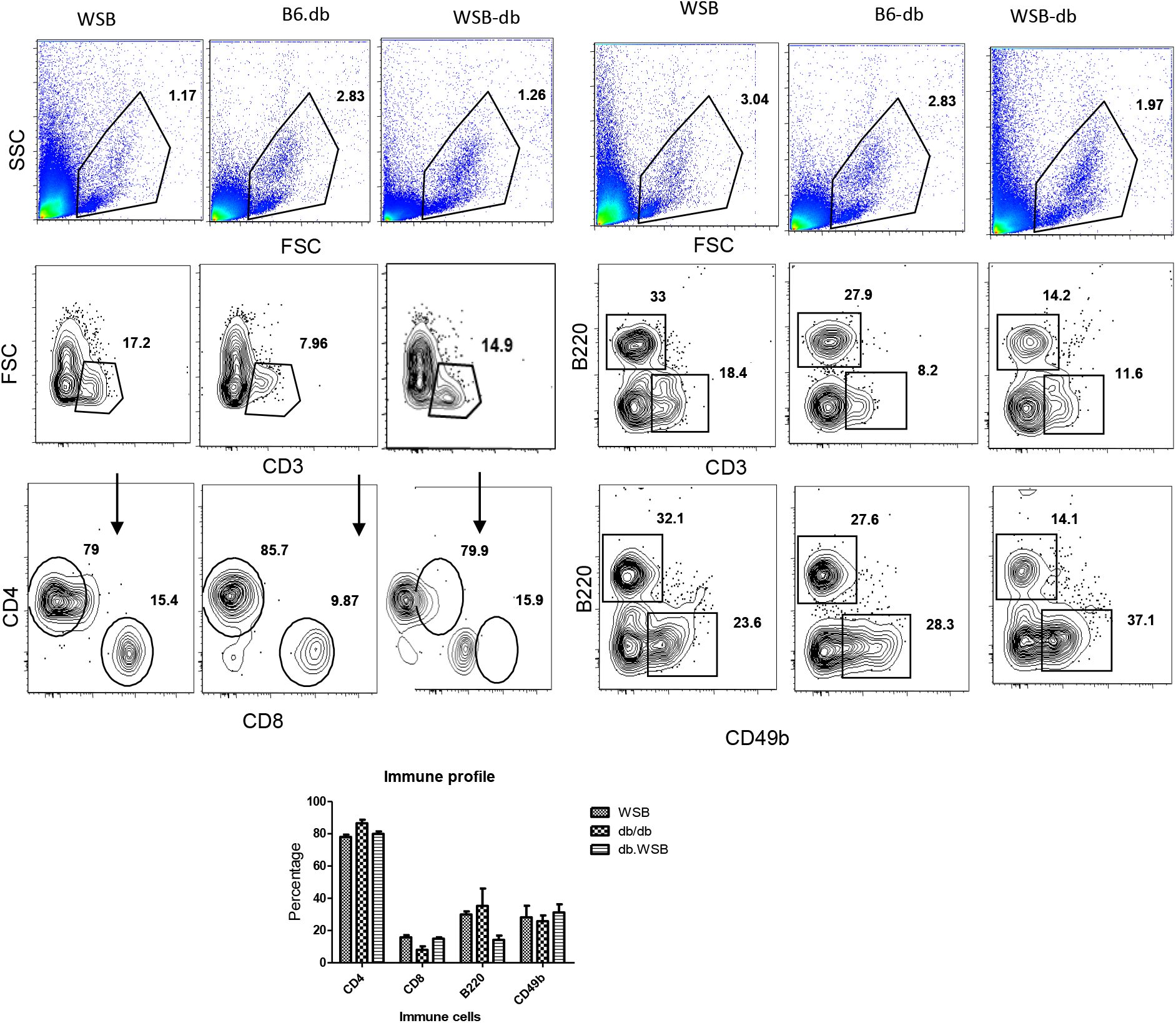
Represents flow cytometer percentage of CD4+, CD8+ B220+ and CD49b population from peripheral blood of WSB,WSB-db,B6-db mice. FACS contour plots of one representative mouse indicating the percentage of CD4+, CD8+ and B220+ and Cd49b cells in blood. Numbers in quadrants indicates percent cells in each throughout. B represents the bar graph of all the positive CD4+, CD8+ and B220+ and CD49a cell population .Statistical analysis was done using 2-way ANOVA. The values are expressed in mean ± SEM. *P < .05, **P < .01, ***P < .001, ****P < .0001

## Discussion

The primary purpose of this study was to understand how differences in mouse genetic background influence obesity and metabolic syndrome. In the present study, we have developed a new congenic strain using WSB as background. The congenic mouse strain (WSB-db) displayed monogenic with moderate obesity pattern that is seen in humans without diabetes and NAFLD and lived longer. Diabetes is under multigenic control and the genome modifiers determine either milder diabetes or diabetes or non-diabetes. Hence the genetic backgrounds of mouse strain play an important role in a disease model. Among the three wild-derived strains, CAST, WSB, PWK, reported WSB mice are resistant to high fat or high sucrose or high fructose for progressing metabolic syndrome ^9^. Extensive studies on metabolic phenotypes in chow and high fat, sucrose-fed WSB mice were studied ^9,10^. Like other wild-derived inbred strains, have found that WSB mice are small and lean as adults, and this strain has been used in studies to genetically map loci affecting body weight and growth rates ^10, 12 and 13.^ Strikingly, male WSB mice are remarkably resistant to the detrimental effects of a diet high in fat and sucrose ^9^. Similarly, others have found male WSB mice are resistant to high fat diet-induced obesity ^14^. Male WSB mice are also resistant to other metabolic effects of high-fat, sucrose feeding. WSB mice do not become hyperglycemic and remain insulin sensitive and glucose tolerant ^9^. The Plasma insulin levels in WSB mice remain low, accompanied by low insulin secretion in vivo ^9^. Considering the above characteristics of WSB/WiJ, we transferred db mutation onto the WSB/EiJ background that results in a profound change in diseases associated with db gene mutation. The food intake in congenic stain (WSB-db) was higher than WSB mice due to *Lepr* mutation that causes hunger with excess food intake, body weight gain, and liver weight gain with adequate white adipose tissue deposition. The above physiologic parameters demonstrated that this strain had developed obesity. On hematology, we found WSB-db mice had a significantly higher lymphocytes count with lower monocytes and granulocytes than WSB and B6-db mice. This may be due to chronic reactive leukocytosis due to an increase in body weight ^15^. Total leukocyte count was associated with liver enzyme levels, insulin resistance as well as visceral and subcutaneous fat thickness. Lymphocyte count was associated with serum liver enzymes, insulin resistance, and dyslipidemia. Monocyte count was associated with serum liver enzyme, insulin resistance, visceral and subcutaneous fat thickness, body fat mass, and percentage body fat ^16^. However, in this study on mice we could not see the correlation except with increase in lymphocytes in WSB-db mice. On biochemistry analysis of serum for liver AST and ALT there was no significant increase and were within the limit of their genotype. However, we could observe significant increase in cholesterol and triglycerides level in WSB-db mice with respect to B6-db and this elevation may be due to increase lipid synthesis. The level of blood glucose, HbA1C was not increased WSB mice as compared to B6.db mice and WSB .db mice indicating a good glycemic control in WSB-mice as reported earlier^10^ the pancreas had reduced growth in WSB mice with increase Insulin expression on Immunofluroscence. However WSB .db mice had marked increase in pancreas size with reducedexpression in beta cells for insulin compared to WSB mice by immunofluorescence. The histopathological evaluation of the liver also corroborated that these congenic mice are relatively resistant to steatosis or any pathological changes in the liver.

The number of literature reports indicates that there are significant alterations in hepatic genes, mRNAs, and proteins expression and functions due to the effect of NAFLD or diabetes. The variants of genes involved in energy metabolism, *denovo lipogenesis*, inflammatory genes and genes involved in glucose metabolism are considered candidates to converse vulnerability to metabolic syndrome, and their upregulation or downregulation effects helps to understand the disease and prevention. In our study, we found that expression of genes involved in energy metabolism, *denovo lipogenesis*, inflammatory genes, and genes controlling glucose metabolism had positive effects in WSB-db regulating the development of NAFLD.

The effects of immune cells changes were also studied in this congenic strain as many studies had been done that absence of Leptin or its receptor results in alterations of the cellularity and function of the immune system. There are studies where Leptin deficiency caused decrease total T cell number, decrease in CD4+ helper T cell number, and a skewing from Th1 to Th2 phenotype protecting certain forms of autoimmunity and increased susceptibility to intracellular infections ^17,18^.

## Conclusion

In summary, the novel WSB-db mouse strain could divulge the existence of modifier genes that can rescue a defective leptin pathway without leptin. The congenic mouse strain developed could serve as an additional model for a genetic and physiological basis for a relatively simple monogenic moderate obesity that is seen in humans with diabetes and resistance to NAFLD, and kidney diseases.

## Acknowledgement

The authors wish to thank the director, National Institute of Immunology, NII and Department of Biotechnology, India for funding support. The authors also wish to thank Mr Subash Dogra SAF for technical help in this study.

